# Pupillary dilation responses as a midlife indicator of risk for Alzheimer’s Disease: Association with Alzheimer’s disease polygenic risk

**DOI:** 10.1101/624767

**Authors:** William S. Kremen, Matthew S. Panizzon, Jeremy A. Elman, Eric L. Granholm, Ole A. Andreassen, Anders M. Dale, Nathan A. Gillespie, Daniel E. Gustavson, Mark W. Logue, Michael J. Lyons, Michael C. Neale, Chandra A. Reynolds, Nathan Whitsel, Carol E. Franz

**Affiliations:** Department of Psychiatry University of California, San Diego, La Jolla, CA, USA; Center for Behavior Genetics of Aging, University of California, San Diego, La Jolla, CA, USA; Center of Excellence for Stress and Mental Health, VA San Diego Healthcare System, La Jolla, CA, USA; Psychology Service, VA San Diego Healthcare System, La Jolla, CA, USA; NORMENT, KG Jebsen Centre for Psychosis Research, Institute of Clinical Medicine, University of Oslo and Division of Mental Health and Addiction, Oslo University Hospital, Oslo, Norway; Department of Radiology, University of California, San Diego, La Jolla, CA, USA; Department of Neurosciences, University of California, San Diego, La Jolla, CA, USA; Virginia Institute for Psychiatric and Behavior Genetics, Virginia Commonwealth University, Richmond, VA, USA; National Center for PTSD: Behavioral Science Division, VA Boston Healthcare System, Boston, MA, USA; Department of Psychiatry and Biomedical Genetics Section, Boston University School of Medicine, Boston, MA, USA; Department of Biostatistics, Boston University School of Public Health, Boston MA, USA; Department of Psychological and Brain Sciences, Boston University, Boston, MA, USA; Department of Psychology, University of California, Riverside, Riverside, CA, USA

## Abstract

Pathological changes in Alzheimer’s disease (AD) begin decades before dementia onset. Because locus coeruleus tau pathology is the earliest occurring AD pathology, targeting indicators of locus coeruleus (dys)function may improve midlife screening for earlier identification of AD risk. Pupillary responses during cognitive tasks are driven by the locus coeruleus and index cognitive effort. Several findings suggest task-associated pupillary response as an early marker of AD risk. Requiring greater effort suggests being closer to one’s compensatory capacity, and adults with mild cognitive impairment (MCI) have greater pupil dilation during digit span tasks than cognitively normal individuals, despite equivalent task performance. Higher AD polygenic risk scores (AD-PRSs) are associated with increased odds of MCI and tau positivity. We hypothesized that AD-PRSs would be associated with pupillary responses in cognitively normal middle-aged adults. We demonstrated that pupillary responses during digit span tasks were heritable (h^2^=.30-.36) in 1119 men ages 56-66. We then examined associations between AD-PRSs and pupillary responses in a cognitively normal subset who all had comparable span capacities (n=539). Higher AD-PRSs were associated with greater pupil dilation/effort in a high (9-digit recall) cognitive load condition; Cohen’s d=.36 for the upper versus lower quartile of the AD-PRS distribution. Results held up after controlling for *APOE* genotype. The results support pupillary response—and by inference, locus coeruleus dysfunction—as a genetically-mediated biomarker of early MCI/AD risk. In some studies, cognition predicted disease progression earlier than biomarkers. Pupillary responses might improve screening and early identification of genetically at-risk individuals even *before* cognitive performance declines.

## INTRODUCTION

Alzheimer’s disease (AD) is a worldwide public health problem and the most expensive disease in the United States^1^. Pathological changes begin decades before onset of dementia, making early identification of AD risk of paramount importance for slowing disease progression^2,3^. Although there are established positron emission tomography and cerebrospinal fluid (CSF) beta-amyloid (Aß) and tau biomarkers, both are costly and invasive. Moreover, several studies have found cognitive function to be an earlier predictor of disease progression than currently defined biomarkers^4–9^. Development of additional, non-invasive markers of risk that might tap some aspect of the disease process even earlier might aid in prediction. We sought to determine if one such marker is a genetically-mediated biomarker of early AD risk.

Postmortem data indicate that tau pathology is the earliest occurring AD biomarker, first appearing in the locus coeruleus (LC)^10–13^. There is also CSF-based evidence indicating that tau pathology can precede Aß in people who progress to AD^14^. Tau is more strongly associated with cognition than Aß^15^, and lower LC neuronal density has been associated with faster cognitive decline in cognitively normal (CN) adults, and individuals with mild cognitive impairment (MCI) and autopsy-confirmed AD^16^. An indicator of LC function may thus be a fitting target to improve screening for earlier identification of AD risk.

Increased pupillary dilation during performance of cognitive tasks is a validated objective psychophysiological index of the brain’s cognitive resource allocation, i.e., cognitive effort^17–20^. Ability level is inversely related to amount of effort—indexed by amount of pupil dilation—needed to perform a task. Pupil size increases with increasing cognitive effort as task demand, i.e., cognitive load, increases^17–21^. However, when task demands substantially exceed abilities and compensatory capacity, there is disengagement from the cognitive processing system; at that point, dilation drops off and performance declines^17–21^. These pupillary responses reflect activation in the LC^22–30^. Although the LC has been viewed historically as important only in terms of broad arousal responses, Aston-Jones and Cohen’s^22^ adaptive gain model supports a complex role of the LC-noradrenergic (LC-NE) system involving phasic activation with adaptive gain to optimize task performance and tonic activation associated with gain that optimizes appropriate disengagement and a shift of focus to different stimuli or tasks. Thus, the LC-NE system is an important modulator of cognitive function and management of cognitive load^22,30–33^.

Taken together, the properties of pupillary dilation responses, their links to LC function, and the potential links between LC tau deposition and development of AD suggest that pupillary dilation responses could anticipate cognitive declines *before* observable performance declines. Suppose two individuals have the same cognitive score. We hypothesized that the one needing more effort is at higher risk for decline because they would be closer to their maximum capacity for compensation^cf.34,35^. On the other hand, someone who has already experienced substantial declines and has surpassed their compensatory threshold is likely to have both poor performance and reduced pupillary dilation responses. Pupillary dilation responses should thus be most useful as a very early marker of risk while there is still little or no observable cognitive decline. Our prior work with participants in the present sample supports these ideas^21^. Individuals with single-domain amnestic MCI had elevated pupillary dilation responses at low or moderate processing loads during digit span tasks, despite equivalent performance to CN participants. Those with multiple-domain MCI had both impaired performance and reduced pupillary responses.

Previously, we showed that a validated AD polygenic risk score (AD-PRS)^36–38^ was associated with increased odds of MCI in participants from the present sample, 89% of whom were <60 years old^39^. The odds ratio for MCI was 3.2 for the upper versus the lower quartile of the PRS distribution. Results changed little after accounting for the effects of *APOE*^39^, the largest single genetic determinant of AD risk^40,41^.

Here we hypothesized that pupillary dilation responses are a genetically-mediated AD risk indicator. We used the classical twin design to estimate the heritability of pupillary responses, thereby demonstrating that they are genetically influenced^42,43^. Next we tested the primary hypothesis that a higher AD-PRS would be associated with greater pupil dilation during a cognitive task even in cognitively normal middle-aged individuals. This association would provide proof of concept supporting the validity and potential utility of pupillary dilation responses as an early marker of risk for MCI and AD.

## MATERIALS AND METHODS

### Participants

Participants were men in wave 2 of the Vietnam Era Twin Study of Aging (VETSA), a national, community-dwelling sample similar to American men in their age range with respect to health and lifestyle characteristics based on Center for Disease Control and Prevention data^44,45^. All served in the military sometime between 1965 and 1975, but ~80% reported no combat exposure. Average age was 61.7 years (SD=2.4; range=56.0-66.9) and average education was 13.8 years (SD=2.1). The average general cognitive ability percentile score was 63.3 (SD=20.7), corresponding to an IQ score of 105^46,47^. Based on a Center for Epidemiologic Studies Depression Scale^48^ threshold of 16, 11.5% met criteria for clinical depression. Also, 27.3% answered yes when asked if they ever had a head injury with loss of consciousness or confusion; almost all were defined as mild and occurred an average of 35 years earlier^49^. Participants traveled to the University of California, San Diego or Boston University where identical protocols were implemented. Written informed consent was obtained from all participants, and the study was approved by Institutional Review Boards at participating institutions.

The present study began with 1207 participants (see Supplementary Figure 1 for flow of participant selection)^44,45^. Exclusions included: self-reported history of glaucoma in either eye, penetrating eye wounds to both eyes, surgery to both eyes involving the muscle, or use of cholinesterase inhibitors or prescribed ocular medications (n=57); or equipment failures or excessive blinking (n=34). Depression and head injury were not exclusions because they are risk factors for dementia. This left 1119 individuals with valid pupillometry data^21^ and 1085 with genotyping data who were of European ancestry. There were too few individuals of non-European ancestry to include in the AD-PRS analyses. There were 828 individuals who were both CN and had valid pupillometry data. [1]Because we were interested in examining whether pupil dilation can inform risk for AD when performance is comparable among individuals, we selected 539 of these 828 individuals with relatively similar maximum span capacities of 5-7 digits (see Discussion for further examination of this issue). Since our digit span task included 3-, 6-, and 9-digit conditions, max span for this subgroup was thus only ±1 digit from the moderate 6-digit load. These included 87 monozygotic (MZ) twin pairs, 62 dizygotic (DZ) twin pairs, and 241 unpaired twins.

### Cognitively Normal Status

As described in detail elsewhere^21,39,50^, cognitive status was determined on the basis of 18 neuropsychological tests covering 6 cognitive domains. Using the Jak-Bondi approach^51^, MCI was defined as having ≥2 tests in a domain that were each >1.5 SDs below normative means. To ensure that MCI reflected a decline in function rather than lifelong low ability, these values were determined after adjusting for general cognitive ability which was assessed at an average age of 20 years^46,52^. Individuals with no impaired domains (85%) were considered CN.

### Pupillometry

We used handheld NeurOptics PLR-2000 pupillometers to record pupil diameter from one eye at 30 Hz for up to 15 seconds while participants viewed a set of lights around a dark interior (~200 lux) inside in a viewing tube. The pupillometer contains recording optics and has a 1.5-inch viewing tube that surrounds the eye and blocks ambient light. To block the other eye, participants closed and held their hand over it. The pupillometer has excellent resolution (mean error=0.052 mm; 99% CI=0.048-0.056; NeurOptics data, N=655).

Pupillary responses were recorded during blocks of trials of 3 (low load), 6 (moderate/near capacity load), and 9 (high/overload) digits presented aurally at the rate of 1 per second. Stimuli were presented on a laptop computer at ~85 decibels. Participants heard “Ready” 1 second before the first digit and “Repeat” 1 second after the last digit. Experimenters initiated pupillary response recording when the word “Ready” was presented. Each trial was inspected for artifacts in a graphic display on the device. Trials were administered until 2 clean trials were recorded or 4 trials were attempted per digit span condition. We averaged trials within each condition and averaged pupil diameter samples for each second of recording (30 per second), corresponding to the presentation of digits at 1-second intervals. The primary dependent variable was the pupillary response score: pupil size at last digit presented minus pupil size at baseline for each trial. These difference scores remove individual differences in tonic pupil size. Supplementary Figure 2 shows a sample pupil response waveform.

### Digit Span Capacity

Maximum span capacity was defined as the longest string of digits correctly recalled during standard tesing with the Wechsler Memory Scale-III digit span subtest without the pupillometer^53^.

### Genotyping Methods

These methods are described in detail elsewhere^39^. Genome-wide genotyping was conducted on individual twins, with one randomly selected twin from each MZ pair at deCODE (Reykjavik, Iceland) with Illumina HumanOmniExpress-24 v1.0A beachips. GenomeStudio software indicated that the average call rate was 0.996. We performed cleaning and quality control with PLINK v1.9^54^. Single nucleotide polymorphisms (SNPs) with >5% missing data or Hardy-Weinberg equilibrium *P*-values<10^−6^ were excluded. Relationships and zygosity were confirmed by PLINKs-genome procedure.

Ancestry was confirmed by SNPweights^55^ and a principal components (PCs) analysis performed in PLINK v1.9 in conjunction with 1000 Genomes Phase 3 reference data^56^. Weights for PCs were computed from 100,000 randomly chosen common (minor allele frequency [MAF]>5%) markers based on 1000 Genomes data and then applied to the VETSA sample. Outliers from the EUR population (1000 Genomes European-ancestry super population) cluster were excluded from the genetically-identified VETSA white non-Hispanic cohort. The remaining white non-Hispanic participants had >89% European ancestry as estimated by SNPweights. PCs for use as covariates to control for potential population substructure within white non-Hispanic participants were recomputed based on 100,000 randomly chosen common markers.

Imputation was performed using MiniMac^57,58^ at the Michigan Imputation Server (https://imputationserver.sph.umich.edu). The 1,000 genomes phase 3 EUR data were used as a haplotype reference panel. Only one randomly chosen individual in each genotyped MZ pair was submitted for imputation. The resulting imputed genotypes were then applied to the co-twin. The final sample with available imputation data included 1,329 individuals.

### AD-PRS Calculation and *APOE* Genotyping

The AD-PRS was computed from summary data of an AD GWAS meta-analysis^41^. It is a weighted average of VETSA sample additive imputed SNP dosages with log-odds ratios for each SNP estimated in the GWAS used as the weights. We excluded rare SNPs (MAF<1%) and SNPs with poor imputation quality (R^2^<0.5) from the calculation. We trimmed the remaining SNPs for linkage disequilibrium (LD) using PLINK’s clumping procedure (r^2^ threshold of 0.2 in a 500 kb window) based on LD patterns in the 1000 Genomes EUR cohort. ADPRSs were computed by PLINK v1.9 using 6 *P*-value thresholds: *P*<0.05, 0.10, 0.20, 0.30, 0.40, 0.50. In addition, we directly genotyped *APOE* as described previously^59^. The number of SNPs included at different thresholds has been documented in a prior publication^39^. In our study of MCI and in studies of AD, the *P*<0.50 threshold provided the best case-control discrimination^36,38,39^. We, therefore, used the *P*<0.50 threshold in the present study.

## Statistical Aanalysis

### Heritability

In the classical twin design, variance of a phenotype is separated into proportions attributed to additive genetic (A), common environmental (C), and unique environmental (E) influences. C influences are environmental factors that make twins in a pair similar to one another; E influences are environmental factors that make twins in a pair different from one another, including measurement error^42,43^. Additive genetic influences are assumed to correlate 1.0 between MZ twins, and 0.50 between DZ twins who on average share 50% of their segregating genes. C influences are assumed to correlate 1.0 between members of a pair regardless of zygosity. E influences are, by definition, uncorrelated between members of a pair. Heritability is the proportion of total variance attributed to additive genetic influences.

Extending to the multivariate case, we examined the relative contribution of the genetic and environmental influences on pupil dilation responses at the 3 cognitive loads and the covariance between these measures by fitting a Cholesky decomposition to the data. The purpose was to determine the degree to which covariance between individual differences at the 3 cognitive loads can be explained by common versus distinct continua of liability. We began by fitting a Cholesky that included the A, C, and E effects, then tested if the A or C components could be removed without any change in model fit. We tested model fit using the likelihood-ratio chi-square test (LRT), which is the difference in the −2 log likelihood (−2LL) of the model in question relative to the full saturated model. Nonsignificant LRT values (*P*>.05) indicate that a reduced model does not have a significantly worse fit relative to the comparison. Additionally, we used the Akaike Information Criterion (AIC) as an indicator of goodness-of-fit; smaller values represent a better balance between goodness-of-fit and parsimony^60^. Analyses were conducted using the raw data option of the maximum-likelihood based structural equation modeling software OpenMx^61,62^.

Residual pupillary response scores were used in the biometrical models, after adjustment for age, pupillometry device (4 of the same devices were used), and medications with anticholinergic properties. Relevant medications and their rankings for degree of anticholinergic properties have been documented previously^21^.

### AD-PRS

These analyses were conducted using linear mixed effects models (SAS Proc Mixed, version 9.4)^63^ accounting for the correlated nature of the twin data by including family (i.e., twin pair) as a random effect. The AD-PRS was standardized prior to analysis. We included the first 3 PCs, age, pupillometry device, and medications with anticholinergic properties as covariates. We also compared the upper versus lower quartile of the AD-PRS distribution. To determine effects of the AD-PRS after accounting for *APOE*, we performed additional analyses including directly genotyped *APOE*-ε2 and *APOE*-ε4. Each was coded for presence/absence of at least one ε2 or ε4 allele, respectively. Results were based on type III tests of fixed effects.

## RESULTS

The full Cholesky provided a good fit to the data (−2LL=4800.15, df=1570, AIC=1660.14). Two C estimates accounted for ≤1% of variance. A reduced Cholesky with those parameters set to zero resulted in minimal change in fit (−2LL=4800.43, df=1575, AIC=1650.43, LRT=.29, df=5, *P*>.999). All 3 pupillary response measures were significantly heritable (h^2^=0.30-0.36); the remaining variance was primarily accounted for by unique environmental influences (Table 1). The unstandardized variance components for the reduced Cholesky also show that the genetic and the total variance in pupillary responses increased as cognitive load increased (Table 1). However, heritabilities changed little with increasing cognitive load because genetic and unique environmental variances were both increasing.

**Table 1.** Variance components of pupillary dilation response measures

| Standardized Variance Components |  |  |  |
| --- | --- | --- | --- |
| Measure | A (95% CI) | C (95% CI) | E (95% CI) |
| *Full Cholesky* |  |  |  |
| Dilation at 3 Digits | .36 (.10;.52) | .00 (.00;.20) | .64 (.48;.83) |
| Dilation at 6 Digits | .33 (.05;.59) | .14 (.00;.38) | .53 (.38;.74) |
| Dilation at 9 Digits | .36 (.06;.54) | .01 (.00;.23) | .63 (.46;.84) |
| *Reduced Cholesky* |  |  |  |
| Dilation at 3 Digits | .36 (.17;.52) | ---- | .64 (.48;.83) |
| Dilation at 6 Digits | .30 (.10;.60) | .17 (.00;.32) | .53 (.38;.73) |
| Dilation at 9 Digits | .37 (.17;.54) | ---- | .63 (.46;.83) |
| Unstandardized Variance Components |  |  |  |
| Measure | A | C | E |
| *Reduced Cholesky* |  |  |  |
| Dilation at 3 Digits | .36 | ---- | .65 |
| Dilation at 6 Digits | .57 | .32 | 1.01 |
| Dilation at 9 Digits | .76 | ---- | 1.26 |

Note: A=Additive genetic influences; C=Common/shared environmental influences; E=Unique environmental influences; CI=Confidence interval.

Table 2 shows the correlations among pupillary response measures derived from the reduced Cholesky. Phenotypic correlations, which represent the total shared variance between measures, were moderate (r_P_=40-0.65). Genetic correlations, which represent only the shared genetic variance between measures^43^, were substantially higher (r_G_=0.73-0.99). The high genetic correlations suggest that genetic influences affecting dilation at varying digit lengths are driven primarily by a single common factor. However, 2 genetic correlations were significantly different from 1.0, indicating that they are not entirely influenced by the same genes. Because unselected samples are thought to provide more unbiased heritability estimates, we also provide the full sample (n=1119) Cholesky and correlation results, which were very similar (Supplementary Tables 1 and 2). However, as already noted, we focus primarily on the subset of individuals with span capacities of 5-7 because of the very different meaning of the task for people at the extremes of span capacity.

**Table 2.** Phenotypic, genetic, and unique environmental correlations among pupillary dilation response measures

| Measures | 3 digits | 6 digits | 9 digits |
| --- | --- | --- | --- |
| *<u>Phenotypic correlations</u>* |  |  |  |
| Dilation at 3 digits | 1.00 |  |  |
| Dilation at 6 digits | .42 (.35 ; .49) | 1.00 |  |
| Dilation at 9 digits | .42 (.35 ; .49) | .60 (.54 ; .65) | 1.00 |
| *<u>Genetic correlations</u>* |  |  |  |
| Dilation at 3 digits | 1.00 |  |  |
| Dilation at 6 digits | .99 (.58 ; 1.0) | 1.00 |  |
| Dilation at 9 digits | .73 (.42 ; .96) | .63 (.18 ; .93) | 1.00 |
| *<u>Unique environmental correlations</u>* |  |  |  |
| Dilation at 3 digits | 1.00 |  |  |
| Dilation at 6 digits | .17 (-.26 ; .36) | 1.00 |  |
| Dilation at 9 digits | .24 (.06 ; .42) | .67 (.53 ; .77) | 1.00 |

Note: Numbers in parentheses are the 95% confidence intervals. All estimates were derived from the reduced trivariate Cholesky decomposition.

The AD-PRS was significantly correlated with pupil dilation response during the 9-digit recall condition (r=0.10, *P*<0.03; Table 3); results for the entire sample were similar, albeit weaker (Supplementary Table 3). The difference between the upper and lower quartiles quartile of the AD-PRS distribution increased as the cognitive load increased (Figure 1). The upper quartile had significantly larger pupil responses during 9-digit recall (Cohen’s d=0.36, *P*<0.005; Table 4), and this comparison was at a trend level for the 6-digit recall (d=0.22, *P*<0.08). These sets of results held up after including maximum span capacity as a covariate, and after controlling for depression and history of head injury (Supplementary Tables 4 and 5). After controlling for presence/absence of directly genotyped *APOE*-ε2 and *APOE*-ε4, the AD-PRS was still significantly correlated with pupil dilation responses during the high cognitive load condition (r=0.11, *p*<0.02; Supplementary Table 6). Neither *APOE* variant was associated with pupil dilation responses.

**Fig. 1.**
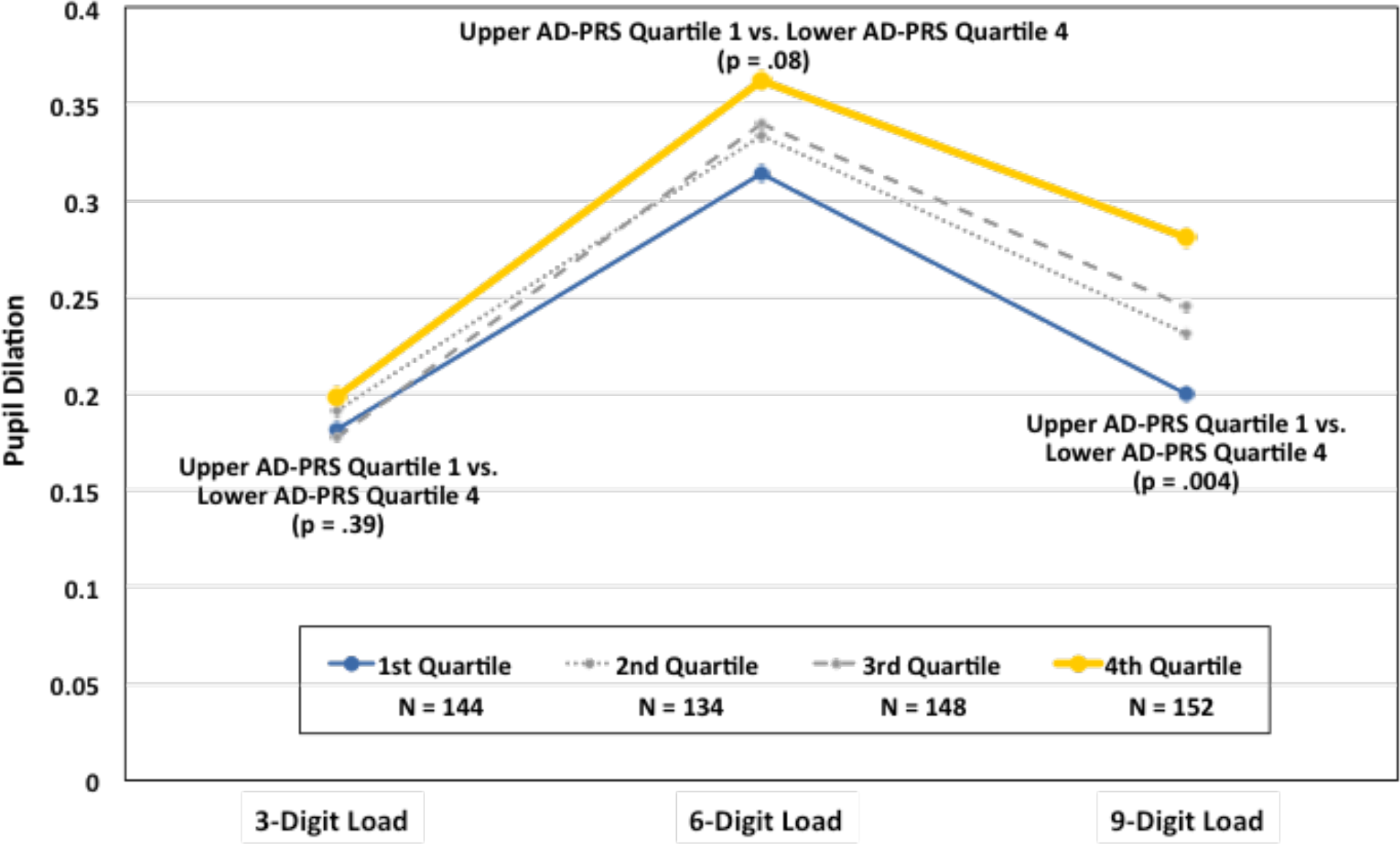
Pupillary dilation response during digit span tasks: Upper vs. lower quartiles of the AD-PRS distribution. AD-PRS = Alzheimer’s disease polygenic risk score.

**Table 3.** Association of Alzheimer’s disease polygenic risk score with pupillary dilation response

| Digit Span Load | Estimate | SE | DF | *t* | *p* | *r* |
| --- | --- | --- | --- | --- | --- | --- |
| 3 Digits (n=537) | 0.003 | 0.007 | 139 | 0.42 | .67 | .02 |
| 6 Digits (n=530) | 0.014 | 0.010 | 135 | 1.42 | .16 | .06 |
| **9 Digits (n=521)** | **0.023** | **0.010** | **130** | **2.18** | **.03** | **.10** |

Note: Covariates include age, the first 3 principal components from the genome-wide genotyping data, pupillometry device, and total number of medications with anticholinergic properties. Data were restricted to cognitively normal individuals with a maximum digit span of 5-7 digits. Ns vary due to missing data for particular variables.

**Table 4.** Association of Alzheimer’s disease polygenic risk score (upper vs. lower quartile) with pupillary dilation response

| Digit Span Load | Estimate | SE | DF | *t* | *p* | *d* |
| --- | --- | --- | --- | --- | --- | --- |
| 3 Digits (n=272) | -0.017 | 0.020 | 137 | -0.86 | .39 | .10 |
| 6 Digits (n=267) | -0.048 | 0.027 | 133 | -1.79 | .08 | .22 |
| **9 Digits (n=264)** | **-0.080** | **0.028** | **128** | **-2.88** | **.005** | **.36** |

Note: Covariates include age, the first 3 principal components from the genome-wide genotyping data; pupillometry device, and total number of medications with anticholinergic properties. Data were restricted to cognitively normal individuals with a maximum digit span of 5-7 digits. Ns vary due to missing data for particular variables. Results presented represent the difference between the upper and lower quartiles of the AD-PRS distribution.

## DISCUSSION

To our knowledge, this is the first evidence of the heritability of task-relevant pupillary dilation responses. High genetic correlations suggest that individual differences in dilation during different cognitive loads are driven primarily by a single common factor or underlying continuum of liability. We then showed that CN individuals at greater genetic risk for AD—based on the AD-PRS—had significantly greater pupil dilation when cognitive demand was high. The effect size comparing the upper and lower quartiles of the AD-PRS distribution was d=.36. Consistent with an underlying continuum of liability There was an increasing effect size with increasing cognitive load,.

We previously observed a wide distribution of pupillary responses in CN individuals, and hypothesized that those with the highest pupil dilation would be at highest risk for progressing to MCI and AD^21^. Although we do not yet know who will develop these disorders, our results support this hypothesis because those who required the greatest effort as cognitive load increased also tend to be those at highest genetic risk based on the AD-PRS. The minimal variation in actual performance in this sample and additional analyses controlling for maximum span show that risk was associated with effort needed rather than task performance. Thus, these results provide proof of concept that pupillary dilation responses during a cognitive task—a brief, low-cost, low-invasive assessment—might be a useful additional risk indicator for identifying participants for clinical trials or other research on determinants of onset and progression of AD.

Although the full sample results were similar, to ensure relatively comparable difficulty level and performance across participants, we only included participants with max spans of 5-7 digits. For individuals with max spans >7, 9 digits is not as much of an overload, and for individuals with a max span of <5 digits, 6 digits is closer to overload. These distinctions are important because, relative to individuals with lower ability, individuals with greater ability dilate less at low loads but more in higher load conditions^19–21^. It is, therefore, important to examine dilation relative to individual ability level.

Here we used pre-set cognitive loads because it was important in our initial work^21^ to show that pupil responses differed in a systematic way as a function of capacity and processing load. Having demonstrated proof of concept, it will be necessary to implement idiographic approaches for meaningful future comparison across all individuals in which cognitive loads are tailored to each individual’s capacity (e.g., defining high load as 2 digits above each individual’s maximum span). Finally, we chose digit span, in part, due to practical constraints of the pupillometry device. However, we have successfully piloted pupil response on a new device with which we can assess episodic memory. Thus, proof of concept demonstrated here will be fully applicable to future studies using idiographic approaches with more AD-relevant episodic memory tests.

Here we acknowledge some limitations. Although this was a community-based sample, it was all male and largely white, non-Hispanic. All had past military service, but the large majority was non-combat-related. Generalization to women or racial/ethnic minorities remains to be determined. We also do not know if the highest cognitive load would best predict risk in other age groups. However, if one’s interest is in biomarkers of early risk for cognitive decline or AD, it is middle-aged adults that may be most appropriate. It will be of interest to determine how AD biomarkers (currently being assessed in this sample) are related to pupillary responses, and if pupillary responses might in some cases detect risk before currently defined Aß and tau thresholds are reached.

### Summary

Pupillary dilation responses are largely driven by the LC-NE system^30,32^, an important modulator of cognitive function^22,31^. The LC is also an early site of tau deposition. This led to our previous work comparing CN and MCI groups, which supports pupillary response as a potential psychophysiological biomarker of risk for MCI and AD^21^. Here we showed that pupillary dilation responses are associated with AD risk genes. Given evidence linking pupillary responses, LC, and tau, the association between the AD-PRS and pupillary response provides additional evidence that is consistent with pupillary responses as a genetically-mediated MCI/AD biomarker. The results provide proof of concept that assessing pupillary responses recorded during cognitive tasks holds promise as a brief, low-cost, low-invasive, first-line screening technique that may aid in identifying adults at increased genetic risk for AD while they are still cognitively normal. Identifying the specific genes associated with the pupillary response factors may improve understanding of the functioning of the LC-NE system and of genetically-mediated factors affecting risk for MCI and AD.

## ACKNOWLEDGMENTS

This work was supported by the National Institute on Aging grants R01AG050595 (W.S.K., M.J.L., C.E.F.), R01AG022381 (W.S.K.), P01AG055367 and R01AG059329 (C.E.F. [sub-PI]), K08 AG047903 (M.S.P.), AG054509 (E.L.G.), Research Council of Norway 223273 (OA.A.), and Stiftelsen KG Jebsen (O.A.A.), KL2TR00144 and P50AG005131 (J.A.E [sub-award]). This material was, in part, the result of work supported with resources of the VA San Diego Center of Excellence for Stress and Mental Health. The Cooperative Studies Program of the U.S. Department of Veterans Affairs also provided financial support for development and maintenance of the Vietnam Era Twin Registry. The content is solely the responsibility of the authors and does not necessarily represent official views of the National Institutes of Health or the Department of Veterans Affairs. We acknowledge the continued cooperation and participation of the members of the VET Registry and their families.

## REFERENCES

1. Alzheimer’s Association. 2018 Alzheimer’s disease facts and figures. Alzheimers Dement 2018; 14(3):367–429.

2. Sperling RA, Aisen PS, Beckett LA, Bennett DA, Craft S, Fagan AM et al. Toward defining the preclinical stages of Alzheimer’s disease: Recommendations from the National Institute on Aging-Alzheimer’s Association workgroups on diagnostic guidelines for Alzheimer’s disease. Alzheimers Dement 2011; 7:280–292.

3. Golde TE, Schneider LS, Koo EH. Anti-abeta therapeutics in Alzheimer’s disease: The need for a paradigm shift. Neuron 2011; 69:203–213.

4. Insel PS, Mattsson N, Mackin RS, Scholl M, Nosheny RL, Tosun D et al. Accelerating rates of cognitive decline and imaging markers associated with beta-amyloid pathology. Neurology 2016; 86(20):1887–1896.

5. Insel PS, Mattsson N, Donohue MC, Mackin RS, Aisen PS, Jack CR, Jr. et al. The transitional association between beta-amyloid pathology and regional brain atrophy. Alzheimers Dement 2015; 11(10):1171–1179.

6. Jansen WJ, Ossenkoppele R, Tijms BM, Fagan AM, Hansson O, Klunk WE et al. Association of cerebral amyloid-beta aggregation with cognitive functioning in persons without dementia. JAMA Psychiatry 2018; 75(1):84–95.

7. Landau SM, Horng A, Jagust WJ, Alzheimer’s Disease Neuroimaging Initiative. Memory decline accompanies subthreshold amyloid accumulation. Neurology 2018; 90(17):e1452–e1460.

8. Jedynak BM, Liu B, Lang A, Gel Y, Prince JL, Alzheimer’s Disease Neuroimaging Initiative. A computational method for computing an Alzheimer’s disease progression score; experiments and validation with the ADNI data set. Neurobiol Aging 2015; 36 Suppl 1:S178–184.

9. Gomar JJ, Bobes-Bascaran MT, Conejero-Goldberg C, Davies P, Goldberg TE, Alzheimer’s Disease Neuroimaging Initiative. Utility of combinations of biomarkers, cognitive markers, and risk factors to predict conversion from mild cognitive impairment to Alzheimer disease in patients in the Alzheimer’s Disease Neuroimaging Initiative. Arch Gen Psychiatry 2011; 68(9):961–969.

10. Duyckaerts C, Braak H, Brion J-P, Buée L, Del Tredici K, Goedert M et al. PART is part of Alzheimer disease. Acta Neuropathol (Berl) 2015; 129(5):749–756.

11. Braak H, Del Tredici K. The pathological process underlying Alzheimer’s disease in individuals under thirty. Acta Neuropathol 2011; 121(2):171–181.

12. Braak H, Del Tredici K. Where, when, and in what form does sporadic Alzheimer’s disease begin? Curr Opin Neurol 2012; 25(6):708–714.

13. Ehrenberg AJ, Nguy AK, Theofilas P, Dunlop S, Suemoto CK, Di Lorenzo Alho AT et al. Quantifying the accretion of hyperphosphorylated tau in the locus coeruleus and dorsal raphe nucleus: The pathological building blocks of early Alzheimer’s disease. Neuropathol Appl Neurobiol 2017; 43(5):393–408.

14. de Leon MJ, Pirraglia E, Osorio RS, Glodzik L, Saint-Louis L, Kim H-J et al. The nonlinear relationship between cerebrospinal fluid Aβ42 and tau in preclinical Alzheimer’s disease. PLOS ONE 2018; 13(2):e0191240.

15. Maass A, Lockhart SN, Harrison TM, Bell RK, Mellinger T, Swinnerton K et al. Entorhinal tau pathology, episodic memory decline, and neurodegeneration in aging. The Journal of Neuroscience 2018; 38(3):530–543.

16. Wilson RS, Nag S, Boyle PA, Hizel LP, Yu L, Buchman AS et al. Neural reserve, neuronal density in the locus ceruleus, and cognitive decline. Neurology 2013; 80(13):1202–1208.

17. Beatty J. Task-evoked pupillary responses, processing load, and the structure of processing resources. Psychol Bull 1982; 91:276–292.

18. Granholm E, Asarnow RF, Sarkin AJ, Dykes KL. Pupillary responses index cognitive resource limitations. Psychophysiology 1996; 33(4):457–461.

19. Ahern S, Beatty J. Pupillary responses during information processing vary with scholastic aptitude test scores. Science 1979; 205:1289–1292.

20. van der Meer E, Beyer R, Horn J, Foth M, Bornemann B, Ries J et al. Resource allocation and fluid intelligence: Insights from pupillometry. Psychophysiology 2010; 47(1):158–169.

21. Granholm EL, Panizzon MS, Elman JA, Jak AJ, Hauger RL, Bondi MW et al. Pupillary responses as a biomarker of early risk for Alzheimer’s disease. J Alzheimers Dis 2017; 56(4):1419–1428.

22. Aston-Jones G, Cohen JD. An integrative theory of locus coeruleus-norepinephrine function: Adaptive gain and optimal performance. Annu Rev Neurosci 2005; 28:403–450.

23. Alnaes D, Sneve MH, Espeseth T, Endestad T, van de Pavert SH, Laeng B. Pupil size signals mental effort deployed during multiple object tracking and predicts brain activity in the dorsal attention network and the locus coeruleus. Journal of Vision 2014; 14(4).

24. Gilzenrat MS, Nieuwenhuis S, Jepma M, Cohen JD. Pupil diameter tracks changes in control state predicted by the adaptive gain theory of locus coeruleus function. Cogn Affect Behav Neurosci 2010; 10(2):252–269.

25. Raizada RD, Poldrack RA. Challenge-driven attention: interacting frontal and brainstem systems. Front Hum Neurosci 2007; 1:3.

26. Rajkowski J, Kubiak P, Aston-Jones G. Correlations between locus coeruleus (LC) neural activity, pupil diameter and behavior in monkey support a role of LC in attention. Society for Neuroscience Abstracts 1993; 19:974.

27. Koss MC. Pupillary dilation as an index of central nervous system alpha 2-adrenoceptor activation. J Pharmacol Methods 1986; 15(1):1–19.

28. Phillips MA, Szabadi E, Bradshaw CM. Comparison of the effects of clonidine and yohimbine on spontaneous pupillary fluctuations in healthy human volunteers. Psychopharmacology (Berl) 2000; 150(1):85–89.

29. Murphy PR, O’Connell RG, O’Sullivan M, Robertson IH, Balsters JH. Pupil diameter covaries with BOLD activity in human locus coeruleus. Hum Brain Mapp 2014; 35(8):4140–4154.

30. Samuels ER, Szabadi E. Functional neuroanatomy of the noradrenergic locus coeruleus: its roles in the regulation of arousal and autonomic function part II: Physiological and pharmacological manipulations and pathological alterations of locus coeruleus activity in humans. Current Neuropharmacology 2008; 6(3):254–285.

31. Sara SJ. The locus coeruleus and noradrenergic modulation of cognition. Nature Reviews Neuroscience 2009; 10(3):211–223.

32. Wilhelm B, Wilhelm H, Lüdtke H. Pupillography: Principles and applications in basic and clinical research. In: Kuhlmann J, Böttcher M (eds). Pupillography: Principles, Methods and Applications. Zuckschwerdt Verlag: München, 1999, pp 1–10.

33. Coull JT, Buchel C, Friston KJ, Frith CD. Noradrenergically mediated plasticity in a human attentional neuronal network. Neuroimage 1999; 10(6):705–715.

34. Riediger M, Li S-C, Lindenberger U. Selection, optimization, and compensation as developmental mechanisms of adaptive resource allocation: Review and preview. In: Birren JE, Schaie KW (eds). Handbook of the Psychology of Aging, 6th edn. Burlington, MA: Amsterdam, 2006, pp 289–313.

35. Stern Y, Arenaza-Urquijo EM, Bartres-Faz D, Belleville S, Cantilon M, Chetelat G et al. Whitepaper: Defining and investigating cognitive reserve, brain reserve, and brain maintenance. Alzheimers Dement 2018.

36. Escott-Price V, Myers AJ, Huentelman M, Hardy J. Polygenic risk score analysis of pathologically confirmed Alzheimer disease. Ann Neurol 2017; 82(2):311–314.

37. Escott-Price V, Shoai M, Pither R, Williams J, Hardy J. Polygenic score prediction captures nearly all common genetic risk for Alzheimer’s disease. Neurobiol Aging 2017; 49:214.e217–214.e211.

38. Escott-Price V, Sims R, Bannister C, Harold D, Vronskaya M, Majounie E et al. Common polygenic variation enhances risk prediction for Alzheimer’s disease. Brain 2015; 138(Pt 12):3673–3684.

39. Logue MW, Panizzon MS, Elman JA, Gillespie NA, Hatton SN, Gustavson DE et al. Use of an Alzheimer’s disease polygenic risk score to identify mild cognitive impairment in adults in their 50s. Mol Psychiatry 2018; 24:421–430.

40. Corder E, Saunders A, Strittmatter W, Schmechel D, Gaskell P, Small G et al. Gene dose of apolipoprotein E type 4 allele and the risk of Alzheimer’s disease in late onset families. Science 1993; 261(5123):921–923.

41. Lambert JC, Ibrahim-Verbaas CA, Harold D, Naj AC, Sims R, Bellenguez C et al. Meta-analysis of 74,046 individuals identifies 11 new susceptibility loci for Alzheimer’s disease. Nat Genet 2013.

42. Eaves LJ, Last KA, Young PA, Martin NG. Model-fitting approaches to the analysis of human behavior. Heredity 1978; 41:249–320.

43. Neale MC, Cardon LR. Methodology for genetic studies of twins and families. Kluwer: Dordrecht, The Netherlands, 1992.

44. Kremen WS, Franz CE, Lyons MJ. VETSA: The Vietnam Era Twin Study of Aging. Twin Res Hum Genet 2013; 16:399–402.

45. Kremen WS, Thompson-Brenner H, Leung YJ, Grant MD, Franz CE, Eisen SA et al. Genes, environment, and time: The Vietnam Era Twin Study of Aging (VETSA). Twin Res Hum Genet 2006; 9:1009–1022.

46. Lyons MJ, Panizzon MS, Liu W, McKenzie R, Bluestone NJ, Grant MD et al. A longitudinal twin study of general cognitive ability over four decades. Dev Psychol 2017; 53(6):1170–1177.

47. Lyons MJ, York TP, Franz CE, Grant MD, Eaves LJ, Jacobson KC et al. Genes determine stability and the environment determines change in cognitive ability during 35 years of adulthood. Psychol Sci 2009; 20:1146–1152.

48. Radloff LS. The CES-D scale: A self-report depression scale for research in the general population. Applied Psychological Measurement 1977; 1:385–401.

49. Kaup AR, Toomey R, Bangen KJ, Delano-Wood L, Yaffe K, Panizzon MS et al. Interactive effect of traumatic brain injury and psychiatric symptoms on cognition among late middle-aged men: Findings from the Vietnam Era Twin Study of Aging. J Neurotrauma 2019; 36(2):338–347.

50. Kremen WS, Jak AJ, Panizzon MS, Spoon KM, Franz CE, Thompson WK et al. Early identification and heritability of mild cognitive impairment. Int J Epidemiol 2014; 43(2):600–610.

51. Jak AJ, Bondi MW, Delano-Wood L, Wierenga C, Corey-Bloom J, Salmon DP et al. Quantification of five neuropsychological approaches to defining mild cognitive impairment. Am J Geriatr Psychiatry 2009; 17:368–375.

52. Genetic determination of cognitive ability during 35 years of adulthood Proceedings of the Cognitive Aging Conference; April 2008; Atlanta, GA.

53. Wechsler D. Wechsler Memory Scale (WMS-III). Psychological Corporation: San Antonio, TX, 1997.

54. Chang CC, Chow CC, Tellier LC, Vattikuti S, Purcell SM, Lee JJ. Second-generation PLINK: Rising to the challenge of larger and richer datasets. Gigascience 2015; 4:7.

55. Chen CY, Pollack S, Hunter DJ, Hirschhorn JN, Kraft P, Price AL. Improved ancestry inference using weights from external reference panels. Bioinformatics 2013; 29(11):1399–1406.

56. 1000 Genomes Project Consortium, Auton A, Brooks LD, Durbin RM, Garrison EP, Kang HM et al. A global reference for human genetic variation. Nature 2015; 526(7571):68–74.

57. Howie B, Fuchsberger C, Stephens M, Marchini J, Abecasis GR. Fast and accurate genotype imputation in genome-wide association studies through pre-phasing. Nat Genet 2012; 44(8):955–959.

58. Fuchsberger C, Abecasis GR, Hinds DA. minimac2: faster genotype imputation. Bioinformatics 2015; 31(5):782–784.

59. Schultz MR, Lyons MJ, Franz CE, Grant MD, Boake C, Jacobson KC et al. Apolipoprotein E genotype and memory in the sixth decade of life. Neurology 2008; 70:1771–1777.

60. Akaike H. Factor analysis and AIC. Psychometrika 1987; 52:317–332.

61. Boker S, Neale MC, Maes H, Wilde M, Spiegel M, Brick T et al. OpenMx: An open source extended structural equation modeling framework. Psychometrika 2011; 76:306–317.

62. Neale MC, Hunter MD, Pritikin JN, Zahery M, Brick TR, Kirkpatrick RM et al. OpenMx 2.0: Extended structural equation and statistical modeling. Psychometrika 2015; 81:535–549.

63. SAS Institute Inc. SAS OnlineDoc 9.4 SAS Institute: Carey, NC, 2013.

