## Supplementary material for "Pupillary dilation responses as a midlife indicator of risk for Alzheimer’s Disease: Association with Alzheimer’s disease polygenic risk": Supplemtary Figure and Tables

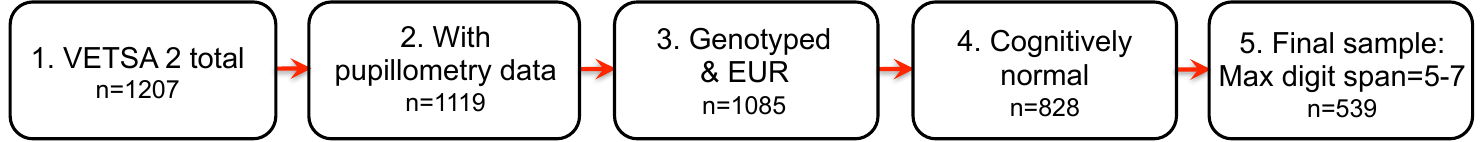

**Supplementary Fig. 1** Flow chart of participant selection.

EUR=Participants of European-American ancestry.

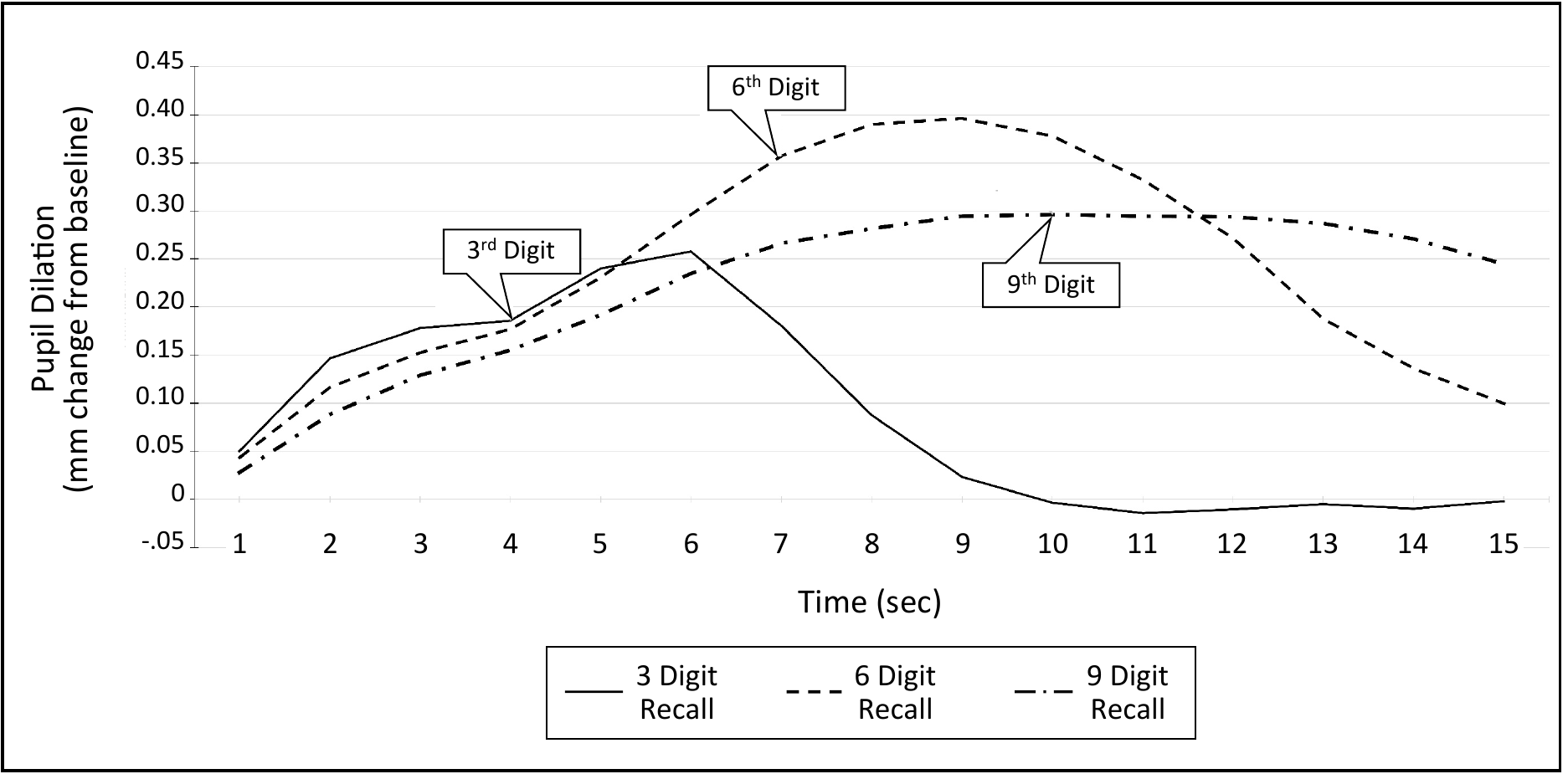

**Supplementary Fig. 2** Sample pupillary response waveform.

| **Supplementary Table 1** Standardized variance components of pupillary dilation response measures in the full VETSA sample (n=1119)^a^ | | | |
| --- | --- | --- | --- |
| Measure | A (95% CI) | C (95% CI) | E (95% CI) |
| *Full Cholesky* |  |  |  |
| Dilation at 3 digits | .25 (.09; .36) | .01 (.00; .15) | .74 (.64; .85) |
| Dilation at 6 digits | .31 (.16; .41) | .00 (.00; .11) | .69 (.59; .80) |
| Dilation at 9 digits | .21 (.05; .36) | .08 (.00; .22) | .71 (.62; .81) |
| *Reduced Cholesky* | |  |  |
| Dilation at 3 digits | .26 (.16; .36) | -- | .74 (.64; .84) |
| Dilation at 6 digits | .31 (.21; .41) | -- | .69 (.59; .80) |
| Dilation at 9 digits | .23 (.12; .37) | .06 (.00; .12) | .71 (.61; .81) |
| A=Additive genetic influences; C=Common/shared environmental influences; E=Unique environmental influences; CI=Confidence interval. All estimates were derived from the reduced trivariate Cholesky decomposition.  ^a^Includes 302 monozygotic twin pairs, 210 dizygotic twin pairs, and 95 unpaired twins. | | | |

| **Supplementary Table 2** Phenotypic, genetic, and unique environmental correlations among pupillary dilation response measures in the full VETSA sample (n=1119)^a^ | | | | |
| --- | --- | --- | --- | --- |
| Measures | 3 digits | 6 digits | | 9 digits |
| *Phenotypic correlations* |  |  | |  |
| Dilation at 3 digits | 1.00 |  | |  |
| Dilation at 6 digits | .47 (.42; .52) | 1.00 | |  |
| Dilation at 9 digits | .40 (.35; .45) | .65 (.61; .68) | | 1.00 |
| *Genetic correlations* |  | |  |  |
| Dilation at 3 digits | 1.00 |  | |  |
| Dilation at 6 digits | .86 (.67; 1.0) | 1.00 | |  |
| Dilation at 9 digits | .73 (.44; 1.0) | .97 (.77; 1.0) | | 1.00 |
| *Unique environmental correlations* | |  | |  |
| Dilation at 3 digits | 1.00 |  | |  |
| Dilation at 6 digits | .31 (.22; .40) | 1.00 | |  |
| Dilation at 9 digits | .31 (.22; .40) | .55 (.48; .62) | | 1.00 |
| Note: Numbers in parentheses are the 95% confidence intervals. All estimates were derived from the reduced trivariate Cholesky decomposition.  ^a^Includes 302 monozygotic twin pairs, 210 dizygotic twin pairs, and 95 unpaired twins. | | | | |

| **Supplementary Table 3** Association of Alzheimer’s disease polygenic risk score with pupillary dilation response in the full VETSA sample | | | | | | |
| --- | --- | --- | --- | --- | --- | --- |
| Digit Span Load | Estimate | SE | DF | *t* | *p* | *r* |
| 3 Digits (n= ) |  |  |  |  |  |  |
| 6 Digits (n= ) |  |  |  |  |  |  |
| **9 Digits (n= )** |  |  |  |  |  |  |
| Note: Covariates include age, the first 3 principal components from the genome-wide genotyping data, pupillometry device, and total number of medications with anticholinergic properties. Data were restricted to cognitively normal individuals with a maximum digit span of 5-7 digits. Ns vary due to missing data for particular variables. | | | | | | |

| **Supplementary Table 4** Association of Alzheimer’s disease polygenic risk score with pupillary dilation response – Controlling for maximum digit span | | | | | | |
| --- | --- | --- | --- | --- | --- | --- |
| Digit Span Load | Estimate | SE | DF | *t* | *p* | *r* |
| 3 Digits (n= ) |  |  |  |  |  |  |
| 6 Digits (n= ) |  |  |  |  |  |  |
| **9 Digits (n= )** |  |  |  |  |  |  |
| Note: Covariates include age, the first 3 principal components from the genome-wide genotyping data, pupillometry device, and total number of medications with anticholinergic properties. Data were restricted to cognitively normal individuals with a maximum digit span of 5-7 digits. Ns vary due to missing data for particular variables. | | | | | | |

| **Supplementary Table 5** Association of Alzheimer’s disease polygenic risk score with pupillary dilation response – Controlling for head injury and depression | | | | | | |
| --- | --- | --- | --- | --- | --- | --- |
| Digit Span Load | Estimate | SE | DF | *t* | *p* | *r* |
| 3 Digits (n=519) | 0.004 | 0.007 | 125 | 0.50 | .62 | .02 |
| 6 Digits (n=513) | 0.016 | 0.010 | 122 | 1.56 | .12 | .07 |
| **9 Digits (n=503)** | **0.022** | **0.011** | **117** | **2.05** | **.04** | **.09** |
| Note: Covariates include age, the first 3 principal components from the genome-wide genotyping data, pupillometry device, and total number of medications with anticholinergic properties. Data were restricted to cognitively normal individuals with a maximum digit span of 5-7 digits. Ns vary due to missing data for particular variables. | | | | | | |

| **Supplementary Table 6** Association of *APOE*-ε2 and *APOE*-ε4 and Alzheimer’s disease polygenic risk score (adjusted for ε2 and ε4) with pupillary dilation response^a^ | | | | | | | | | | | | | | | |
| --- | --- | --- | --- | --- | --- | --- | --- | --- | --- | --- | --- | --- | --- | --- | --- |
|  | *APOE*-ε2 | | | |  | *APOE*-ε4 | | | |  | AD-PRS (adjusted for ε2 & ε4) | | | | |
|  | Estimate | SE | t | p |  | Estimate | SE | t | p |  | Estimate | SE | t | p | r |
| 3-Digit Load (n=526) | 0.022 | 0.020 | 1.13 | .26 |  | 0.006 | 0.016 | 0.40 | .69 |  | 0.004 | 0.007 | 0.50 | .62 | .02 |
| 6-Digit Load (n=519) | -0.006 | 0.028 | -0.23 | .82 |  | -0.019 | 0.022 | -0.89 | .38 |  | 0.015 | 0.010 | 1.45 | .15 | .06 |
| **9-Digit Load**  **(n=510)** | -0.010 | 0.029 | -0.33 | .74 |  | 0.005 | 0.023 | 0.21 | .83 |  | **0.025** | **0.011** | **2.42** | **.02** | **.11** |
| Note: Covariates include age, the first 3 principal components from the genome-wide genotyping data; pupillometry device, total number of medications with anticholinergic properties, and *APOE*-ε2 and *APOE*-ε4 status. Data were restricted to cognitively normal individuals with a maximum digit span of 5-7 digits.  ^a^*APOE*-ε2 and *APOE*-ε4 status were based on direct genotyping. Each was coded as present versus absent for at least one ε2 or ε4 alleles, respectively. | | | | | | | | | | | | | | | |
